# An analytical derivation of the distribution of distances between heterozygous sites in diploid species to efficiently infer demographic history

**DOI:** 10.1101/2023.09.20.558510

**Authors:** Peter F. Arndt, Florian Massip, Michael Sheinman

## Abstract

Heterozygous sites are not uniformly distributed along a diploid genome. Rather, their density varies as a result of recombination events, and their local density reflects the time to the last common ancestor of the maternal and paternal copies of a genomic region. The distribution of the density of heterozygous sites therefore carries information about the history of the population size. Despite previous efforts, an exact derivation of the distribution of heterozygous sites is still lacking. As a consequence, the estimation of population size variation is difficult and requires several simplifying assumptions. Using a novel theoretical framework, we are able to derive an analytical formula for the distribution of distances between heterozygous sites. Our theory can account for arbitrary demographic histories, including bottlenecks. In the case of a constant population size the distribution follows a simple function and exhibits a power-law tail proportional to *r*^*α*^ with *α* =−3, where *r* is the distance between heterozygous sites. This prediction is accurately validated when considering heterozygous sites in individuals of African descent. Other populations migrated out of Africa and underwent at least one bottleneck which left a distinct mark on their interval distribution between heterozygous sites, i.e., an overrepresentation of intervals between 10 and 100 kbp in length. Our analytical theory for non-constant population sizes reproduces this behavior and can be used to study historical changes in population size with high accuracy. The simplicity of our approach facilitates the analysis of demographic histories for diploid species, requiring only a single unphased genome.

## 1 Introduction

The evolution of genomes is driven by the interplay of several evolutionary forces. Mutagenesis initially introduces genomic variation into the genome of a single individual, which is then subject to natural selection and genetic drift. The relative strength of these two mechanisms depends on the selective advantage or disadvantage conferred by a mutation and the size of the population. In large populations, genetic drift is weak and natural selection is the dominant evolutionary force, whereas in small populations, genetic drift is strong and natural selection is weak. Therefore, demographic events such as population bottlenecks can have a strong effect on genome evolution and leave characteristic traces in genomes.

In diploid species that reproduce sexually, genetic recombination is another important force shaping genome evolution. In these species, chromosomes come in pairs, one inherited from the father and one from the mother. However, as a result of crossing over events during meiosis, chromosomes recombine, and a single gamete passed on to the next generation carries a combination of maternal and paternal genetic information (see Fig. 1(A)). Thus, over several generations, each of an individual’s two sister chromosomes is a mosaic of segments of genetic material from its ancestors.

**Figure 1:**
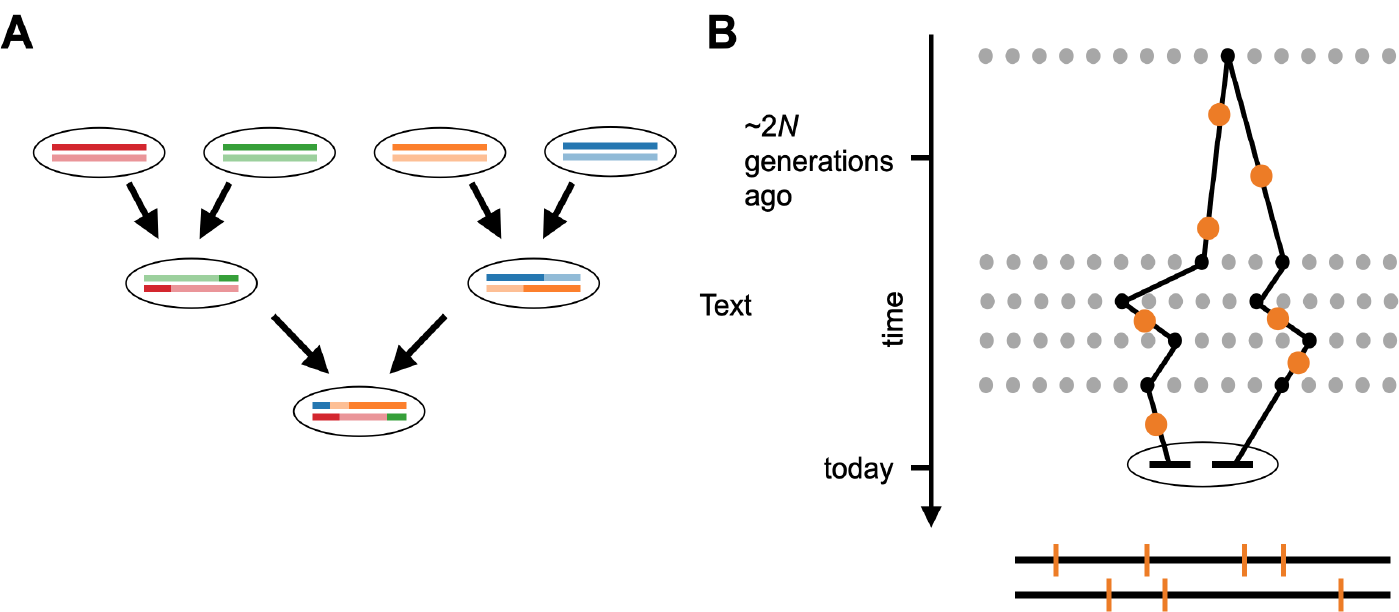
(A) The mosaic structure of ancestral genomic information in a sexually reproducing diploid individual. Each line represents a sister chromosome. Due to crossover during meiosis, a combination of maternal and paternal genomic material is passed on to the next generation. In a finite population, each segment will coalesce at a segment-specific TMCRA. (B) The evolutionary history of an IBD segment. The black line traces the maternal and paternal lineage through ancestral gametes (represented as gray dots) back to the most recent common ancestor. At the TMRCA both copies are identical and accumulate mutations (indicated by orange dots). In the shown example the 4 + 3 mutations along the two lineages break the IBD segment into 8 IBS tracts as indicated at the bottom.

In a finite population, however, the ancestral lineages of each such segment will eventually coalesce at the time of the most recent common ancestor (TMRCA), see Fig. 1(B). These segments are therefore said to be *identical by descent* (IBD). At the time point of coalescence, these segments were also *identical by state* (IBS), i.e. their genomic sequence was 100% identical. Subsequently, both copies accumulated mutations over evolutionary timescales. These mutations are detected as heterozygous sites in a diploid genome that break IBD segments into shorter IBS tracts that are themselves homozygous, see Fig. 1(B). Even if one assumes that these mutations occur uniformly along individual IBD segments, on longer length scales the density of heterozygous sites along chromosomes becomes non-uniform, due to the fact that different IBD segments have different TMRCAs. In fact, it has been observed that the density of single nucleotide polymorphisms (SNPs) varies along genomes by orders of magnitude, much more than expected for a random Poisson process. [1, 2, 3, 4].

In the past, the density of SNPs along chromosomes was already analyzed to infer properties of the coalescent process, in particular the effective population size. Several methods based on the Sequential Markovian Coalescent (SMC) have been developed to infer the demographic history of a genomic sample from SNP data. These methods have in common that they model the Ancestral Recombination Graph, i.e. the genealogies of the observed sample along the genome, assuming that the genealogies follow a Markov process [5]. The first such method, the pairwise sequentially Markovian coalescent (PSMC) [6], was used to infer demographic history from a single diploid genome. Later, this method was extended to analyze multiple pairs of haploid genomes from a population in a method known as multiple sequentially Markovian coalescent (MSMC) [7]. To enable the analysis of large population samples of hundreds of individuals, a novel regularization scheme that reduces estimation error was introduced with SMC++ [8]. Many other extensions [9, 10, 11, 12] have been proposed to address the limitations of SMC-based algorithms for inferring the demographic history of populations (see [13] for a review). In parallel, a competing approach to infer the demographic history was proposed by Harris and Nielsen [14]. They proposed to examine the length distribution of IBS tracts and derived an approximate closed-form formula for the expected length distribution of IBS tracts under different demographic scenarios. However, due to the mathematical complexity of their method, it is rarely used compared to PSMC or similar approaches.

Related to the density of heterozygous sites is the length of homozygous segments. Historically, and due to the fact that long Runs of Homozygosity (RoH) have been associated with inbreeding and genetic diseases, researchers first used restriction fragment length polymorphisms to identify long RoH and map disease loci [15]. Data from about 8, 000 short tandem repeat polymorphisms [16] and later high-density genome-wide scans using SNP microarrays interrogating 3 million SNPs [17] allowed RoH to be found at higher resolutions (up to the 1 kbp scale) and showed that RoH are common in human populations [18]. Finally, short-read whole genome sequencing now surpasses microarray scans in resolution and precision, and phased genotype data are available for more than 1000 individuals in several human populations [19].

The aim of this article is to develop a mathematical model for the length distribution of RoH or IBS tracts. We propose to revisit the approach of Harris and Nielsen [14] and show for the first time that the length distribution of IBS tracts follows a simple function and exhibits a power-law tail *r*^*α*^ with exponent *α* = −3, where *r* is the length of an IBS tract. Here we provide a simple theoretical framework for computing this distribution. We test our theory using empirical data from the 1000 Genomes Project [19, 20] and find that it agrees very well with empirical data for humans of African ancestry. We also derive an extension of our model for individuals who have undergone a population bottleneck, and show that we can use this extended model to date and estimate the magnitude of population bottlenecks.

## 2 Results

The first type of question we want to answer concerns the number and distribution of heterozygous sites in a genome. The number of heterozygous sites of an individual provides valuable information about the genetic diversity in a population. A higher number of heterozygous sites indicates greater genetic diversity within a population. The total number of heterozygous sites in a diploid genome of length *L* is well approximated by *Lθ* with the scaled mutation rate *θ* = 4*N*_*e*_*μ*, the mutation rate per bp and generation, *μ*, and the effective size of a panmictic population in equilibrium without selection, *N*_*e*_ [21, 22, 23, 24].

For humans we have *θ* ≪ 1 and expect a heterozygous site every

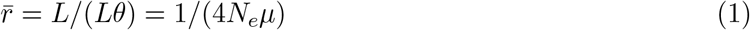

nucleotides along the genome on average. However, it is known that the local density of heterozygous sites along the chromosomes is not uniform [1], see Fig. 2(A), Very similar density distributions can also be observed when considering SNPs rather than heterozygous sites. Note, however, that we will not consider homozygous SNPs in an individual in the following as their presence depends on the underlying reference genome.

**Figure 2:**
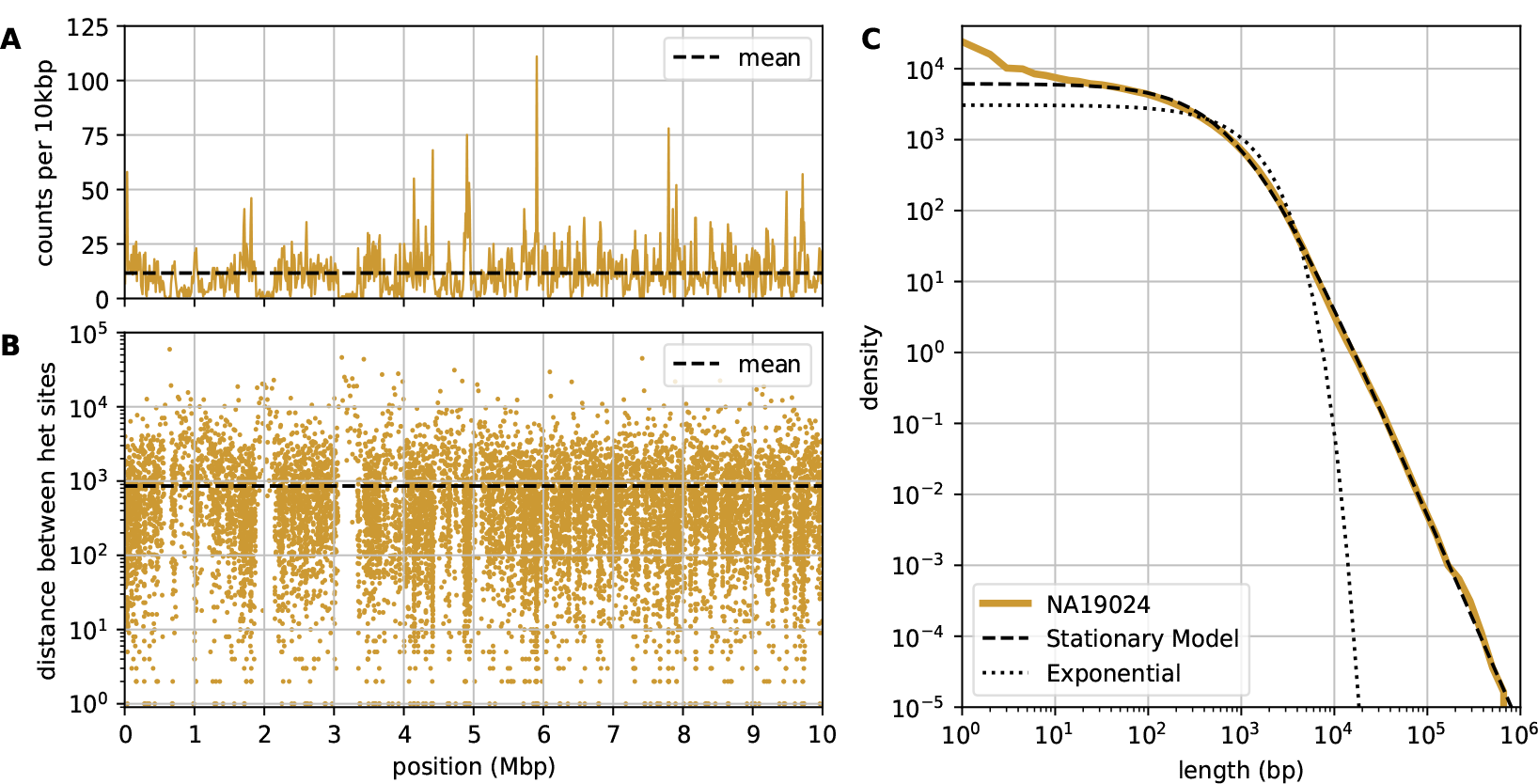
(A) A density profile of heterozygous sites along a 10 Mbp region on chromosome 1 of an individual (NA19024) of African ancestry. Counts are in windows of 10 kbp, the mean count (11.6) is indicated by the dashed line. (B) the same data in a rainfall plot showing the logarithm of the distance between consecutive heterozygous sites against their position along the chromosome. The mean distance (857 bp) is indicated as a dashed line. These distributions are remarkably different from randomly shuffled heterozygous sites in Fig. S1. (C) The genome wide length distribution of IBS tracts of this individual. Shown is the density of counts in logarithmic bins along the *x*-axis. The dotted line represents an exponential distribution with the same mean and the the dashed line represents a fit with our stationary model. The fitted value for *L* = 2.7 *·* 10^9^ and *θ* = 1.06066 *·* 10^−3^. The slope of the tail is −3 corresponding to a power-law *∝ r*^−3^.

To get a better understanding of the density distribution of heterozygous sites we consider the same data in a rainfall plot [25], see Fig. 2(B). In this plot we clearly see that genomes are a mosaic of regions with various densities of heterozygous sites. Each “piece” of this mosaic is an IBD segment which dates back to a different most recent common ancestors and therefore had the potential to accumulate more or less mutations [1, 14].

Here we want to focus on the distribution of distances between heterozygous sites or the length distribution of IBS tracts. This distribution for an individual of African ancestry is shown in Fig. 2 (C). and is very different from the distribution one would obtain if heterozygous sites were uniformly distributed along the genome (see Fig. S1). Such a distribution of long IBS tracts was also observed previously [14]. Here we find that this distribution has a specific shape and exhibits a power-law tail *M* (*r*) ≃*r*^*α*^ with *α* = −3. This intriguing observation can be understood by a mathematical model with only a few simplifying assumptions as deduced below.

Heterozygous sites of an individual genome represent sites where the maternal and paternal alleles differ. Each IBD segment has a different time to the most recent common ancestor (TMRCA). For an individual genome, the TMRCA distribution depends on the structure of the population. For a stationary population with *N* diploid individuals, we have that the distribution of TMRCAs, is given by the probability of coalescence:

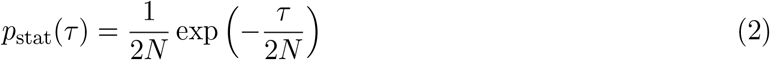

where *τ* is the TMRCA [24, 26]. For segment with TMRCA *τ* the length distribution of IBS tracts is given by a stick-breaking process [27, 28] and follows

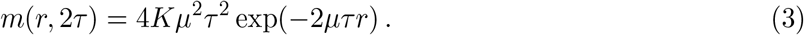

where *r* is the length of a IBS tract, *μ* the mutation rate, and *K* the length of the IBD segment, which is assumed to be larger than a typical IBS tract. Combining Eqs. (2) and (3), assuming that recombination is acting on length scales longer than a typical IBS tract it follows that the genome-wide length distribution of IBS tracts *M* is

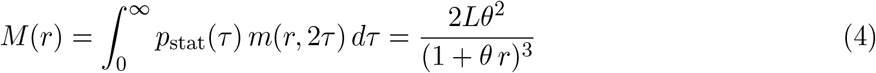

where *L* is the length of the considered genome. This distribution has a mean value 1*/θ* and shows the characteristic power-law tail *∝ r*^*α*^ with exponent *α* = −3 as observed in Fig. 2 (C). Beside the sequence length *L*, the model has only one more free parameter *θ* and the observed empirical distribution for individuals of African ancestry can be very well fitted by our model, see Fig. 2(C). Note that the power-law exponent *α* does not depend on any model parameter. Assuming that the mutation rate *μ* = 2.36*·*10^−8^ per generation and bp [12], the fitted value for *θ* = 0.00106066 corresponds to a effective population size of *N*_0_ = 11, 280 *±* 80.

However, considering individuals with different ancestry, especially those which are assumed to have underwent a population bottleneck while moving out of Africa, we observe significant deviations from our predictions (Fig. 3(C-F) and Fig. 5). Notably, we observe an up to twofold excess of IBS tracts of lengths between 10^4^ and 10^5^ bp. Such deviations were previously shown to be the result of bottleneck events [14].

**Figure 3:**
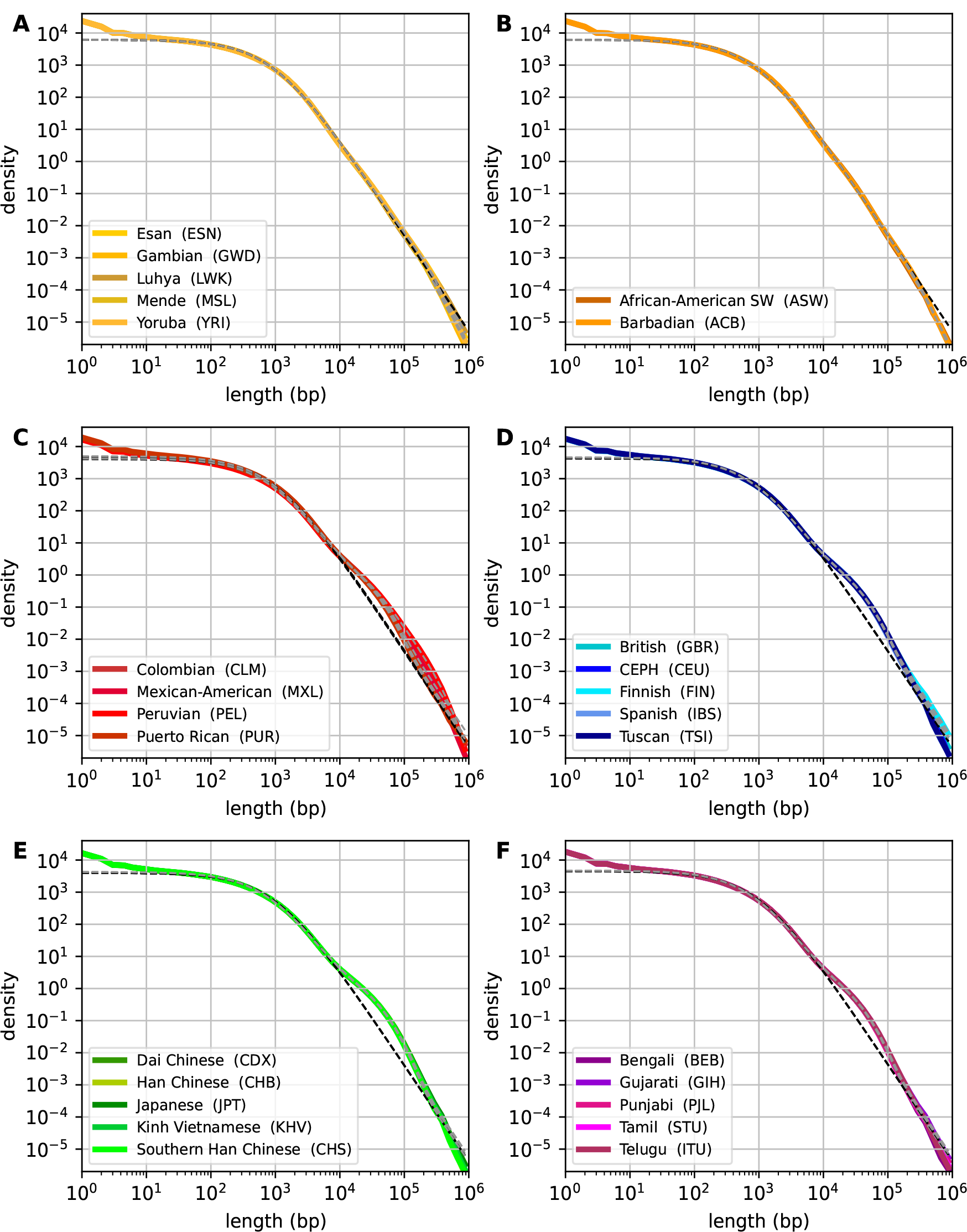
The length distributions of IBS tracts for individuals of (A) African, (B) African American (C) South American (D) European (E) East Asian, and (F) South Asian ancestry. We present the aggregated length distributions over all individuals in an ancestry group. The empirical data is fitted using the stationary model and the bottleneck model, as shown in black and gray, respectively.

To calculate analytically how demographic events such as bottlenecks affect the IBS tract length distribution, we first compute statistical properties of coalescent times of IBD segments in population with varying population size.

Let us consider a population of diploid individuals evolving in time *t* with a varying population size *N* (*t*). Each individual diploid genome is partitioned into IBD segments, i.e. genomic regions for which all sites of the maternal and paternal haploid copy coalesce at the same most recent common ancestor, which we denote by *τ* . In a population of size *N* there are 2*N* haploid genomes, and (2*N*)^2^*/*2 possible pairs of haploid genomes. We define *n*(*τ, t*) as the number of IBD segments with TMRCA *τ* in all pairs of haploid genomes in the population normalized by the length of the genome *L* at time *t*. In other words the expected probability to coalesce at TMRCA *τ* for two homologous base pairs in the population is *n*(*τ, t*)*/*2*N* ^2^. Extending previous considerations [29, 30] we find that the time evolution of *n*(*τ, t*) is given by

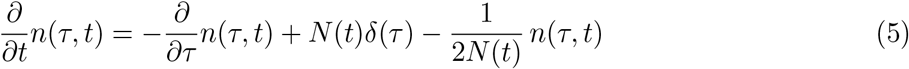

where the three terms on the r.h.s. describe contributions to the evolution of *n*(*τ*) in time due to *(i)* pre-exiting pairings with finite TMRCA *τ, (ii)* new pairings with *τ* = 0 appearing due to coalescent events, and *(iii)* the loss of pairings due to the death of individuals. Note that *N* (*t*) might vary in time and that the normalisation of *n*(*τ, t*) is such that the total number of pairings at each time equals

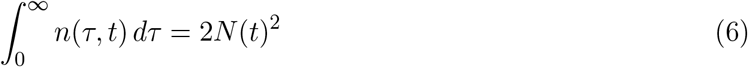

which is the total number of possible pairs assuming that *N* (*t*) *≫* 1. Considering the Laplace transform of *n*(*τ, t*) in *τ* :

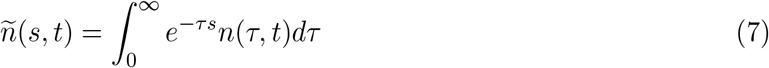

the differential Eq. (5) takes the form

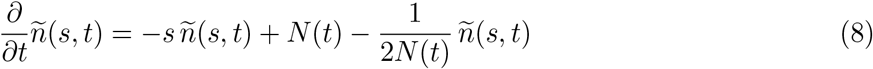

and can be solved by integration. From a solution *n*(*τ, t*) one can compute the length distribution of IBS tracts in a genome of size *L* using the stick breaking process with stick length distribution (3):

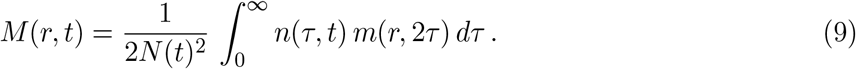

Due to the relationship of the integral in Eq. (9) to Laplace transformations, we can also compute the length distribution of IBS tracts directly as the second derivative of the Laplace transform of *n*:

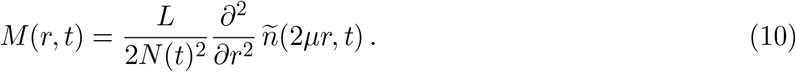

With these computational simplifications we are now in the position to compute the length distribution of IBS tracts in closed form.

As an example consider again a stationary population of constant size *N*. The number of pairs and its Laplace transform *ñ*-(*τ, t*) do not depend on time, i.e. *∂ñ*(*s, t*)*/∂t* = 0, and we solve Eq. (8) by

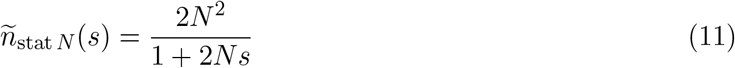

Using Eq.(10) the length distribution of IBS tracts is therefore

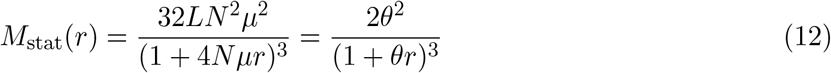

as found above using simpler arguments leading to Eq. (4).

Let us now consider a scenario for which the population size is piece-wise constant for given time intervals, see Fig. 4. We start with a stationary population of size *N*_0_. At time *t* = 0 this population expands or shrinks to size *N*_1_ for a duration of time *T*_1_. Subsequently the population size changes to *N*_*i*_ for a time *T*_*i*_ for of total *E* epochs *i* = 1, 2,…, *E*. We define *t*_*k*_ =Σ_*i*≤k_ *T*_*i*_ as the time points of population size changes. Finally, we observe and analyse the genome of an individual in a population of size *N*_*E*_ at time *t*_*E*_.

**Figure 4:**
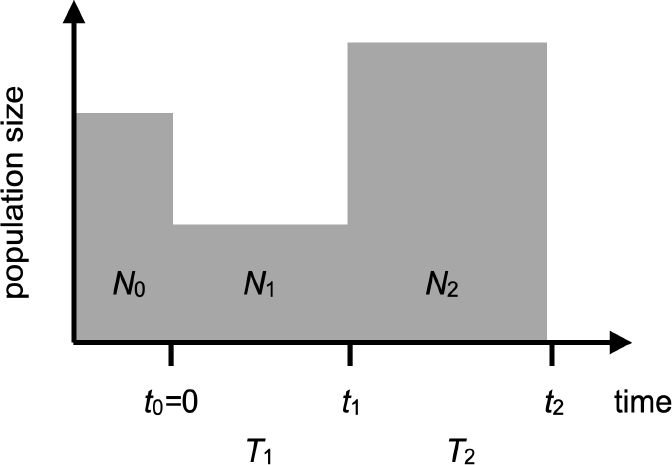
A sketch for the population size in the bottleneck model with *E* = 2 epochs. The initial stationary population is of size *N*_0_. At time *t*_0_ = 0 the population size is reduced to *N*_1_ for time *T*_1_ and then increases to *N*_2_ for time *T*_2_. We observe individuals at time *t*_2_ = *T*_1_ + *T*_2_.

**Figure 5:**
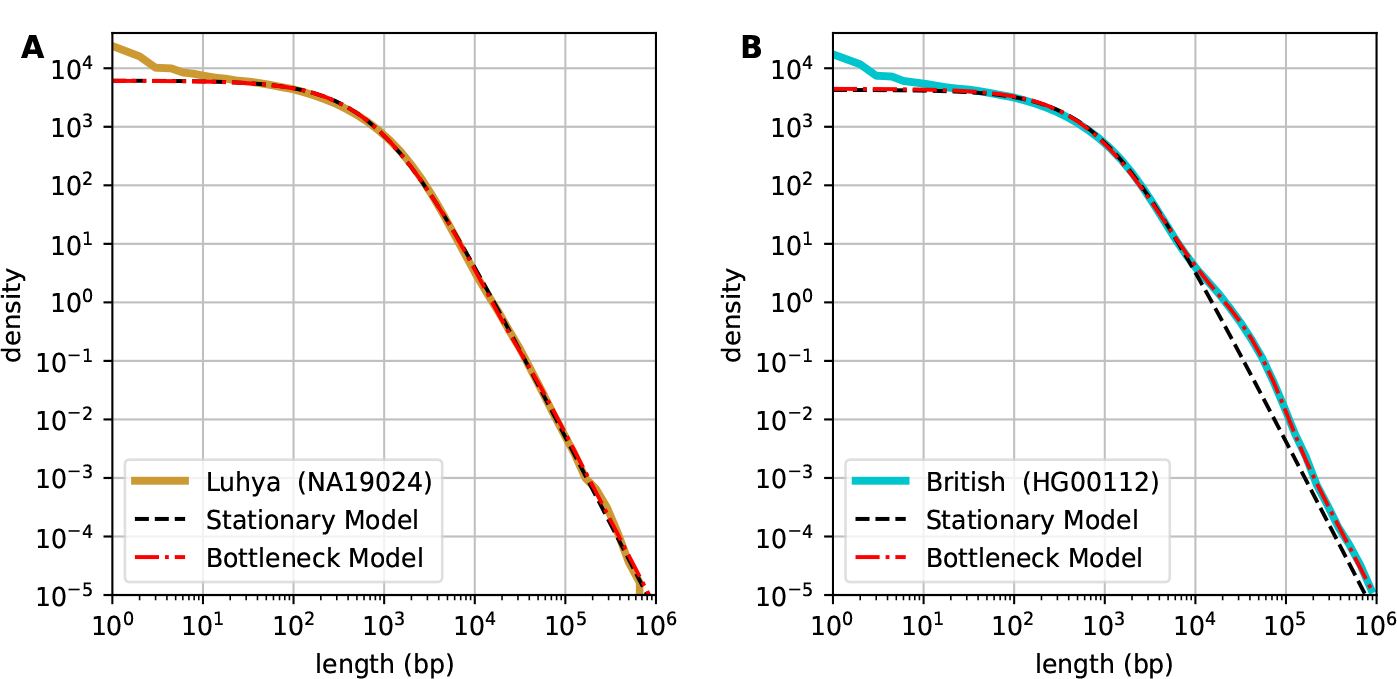
The length distribution of IBS tracks *M* (*r*) of an (A) African and (B) European individual together with fits of two different models.

The solution of the differential Eq. (8) can be computed iteratively by considering an initially stationary population at time *t*_0_ = 0 with

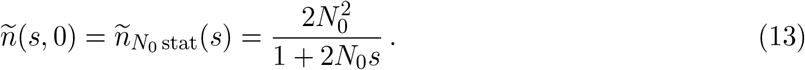

At the time points *t*_*i*_ for *i* = 0, 1,…,*E* − 1 the population of size *N*_*i*_ with pairings *ñ*(*s, t*_*i*_) is subjected to either a reduction or an expansion of the population size.

In case of *(i)* an **instantaneous reduction of the population size**, i.e. if *N*_*i*+1_ ≤ *N*_*i*_ we just take a sample of size *N*_*i*+1_ individuals. This changes the normalisation of the function *n*, i.e. at time point 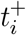 just after the population size change:

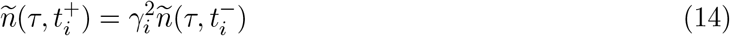

where *γ*_*i*_ = *N*_*i*+1_*/N*_*i*_ ≤ 1 and 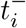 denotes a time just before *t*_*i*_.

In contrast, in case of *(ii)* an **instantaneous expansion of the population size**, i.e. *N*_*i*+1_ *> N*_*i*_, we model that each gamete will generate several offspring to make up the next generation. We assume that the number of offspring, *p*, for each gamete is distributed according to a truncated Poisson distribution:

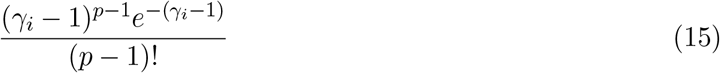

for *p≥*1 and vanishes for *p* = 0, i.e. we assume and that no gamete is lost. On average each gamete will have *γ*_*i*_ = *N*_*i*+1_*/N*_*i*_ *>* 1 offspring and the resulting population at time 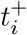 will be of size *N*_*i*+1_. The number of pairings in Laplace space is

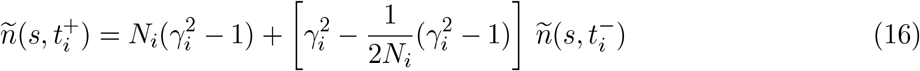

which is a simple affine transformation of the number of pairings before the population size increase. In sum, both a decrease and an increase of the population size amounts to an affine transformation of *ñ*(*s, t*) in the following denoted by 𝒜_*Ni,Ni*+1_. Note that if the population size does not change,*N*_*i*_ = *N*_*i*+1_ the solution is consistent and we have 𝒜_*N,N*_ = id.

Between consecutive changes of the population size, during the time interval *T*_*i*_, we need to **prop-agate** the solution 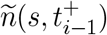 to 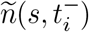 with constant population size *N*_*i*_. Using Eq. (8) we have

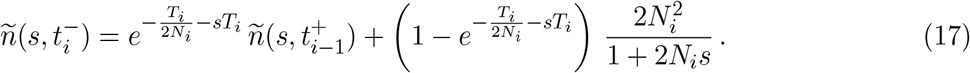

We denote this transformation by 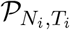.

In summary, by combining these three simple transformations of the initially stationary distribution 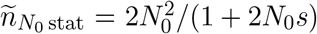 at *t*_0_ = 0 we can iteratively compute the distribution *ñ* at time *t*_*E*_ under the influence of the intermediate population size changes as

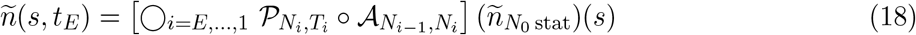

In the end the length distribution of IBS tracts *M* is computed as the second derivative of the final *ñ*(*s, t*_*E*_) following Eq. (10).

Our result above allow us to efficiently compute the length distribution of IBS tracts *M* for relevant evolutionary scenarios. We find that the following model can accurately describe the observed length distribution of IBS tracts for populations that migrated out of Africa. We consider a model with *E* = 2 epochs: an initial population with *N*_0_ individuals goes through a bottleneck such that its population size shrinks to *N*_1_ for a time *T*_1_ and then grows again to a population of size *N*_2_ for a time *T*_2_. With this model we obtain very accurate fits of IBS tracts distributions for populations with different ancestries, see Fig. 3 where we combined IBS tracts for all individuals in an ancestry group (see Methods). Importantly, our model convincingly recovers the overrepresentation of long IBS tracts in populations that have migrated out of Africa, and can also be applied on single individuals, see Figs. 5(A) and 5(B) for exemplary individuals of African and European ancestries.

Comparing the two models (with and without a bottleneck) for an individual of African ancestry, the one with a bottleneck shows no significant improvement (see Fig. 5(A)), although the bottleneck model has 4 more parameters than the model with a stationary population. We further consider bootstrapped samples of the length distributions to compare the two models. For an individual of European ancestry we observe a significant increase (p *<* 10^−20^) of the likelihoods for the model with a bottleneck, Fig. 6. In contrast, the likelihood distributions for fits for an African individual are overlapping indicating that the bottleneck model for African populations is not justified.

**Figure 6:**
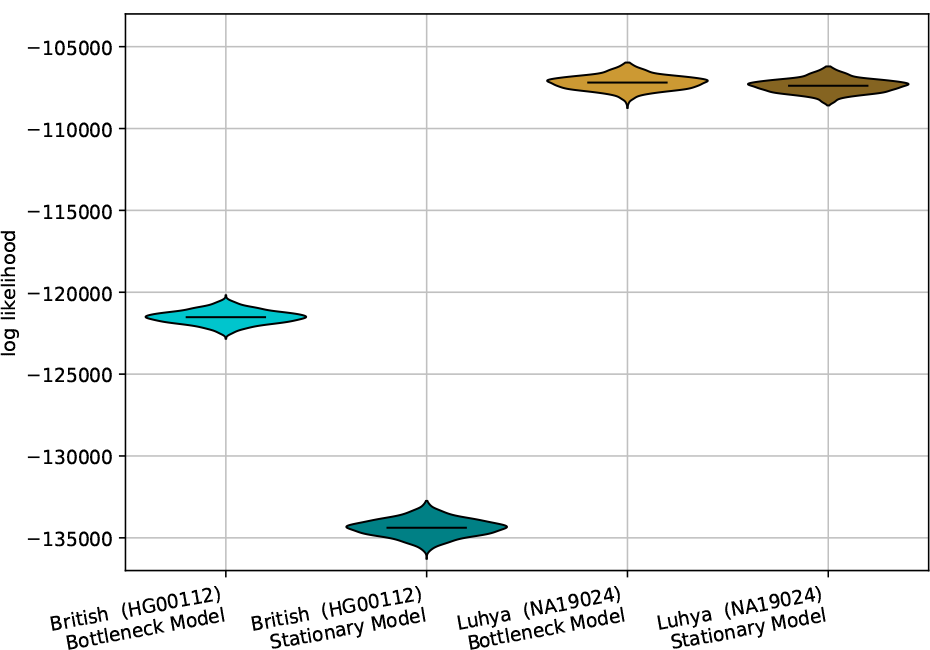
Log likelihood distributions of two model fits to re-sampled length distributions of IBS tracts of two individuals, one of European and one of African ancestry. The model including a population bottleneck significantly improves the model fit for the European individual.

Next we fitted the stationary population model and the *E* = 2 epochs bottleneck model to all individuals in the 1000 genomes dataset [19]. As observed in the example (see Fig. 5(B)), we see significant improvements of the fits using the bottleneck model for South American, European, and Asian populations relative to the stationary population model, Fig. S2. The fitted demographic parameters are shown in Fig. 7. The estimates for *N*_0_ are all very consistent within a 2% relative error margin and point to an ancestral effective population size of about 11, 280 *±* 80 individuals, see also Tab. S1.

**Figure 7:**
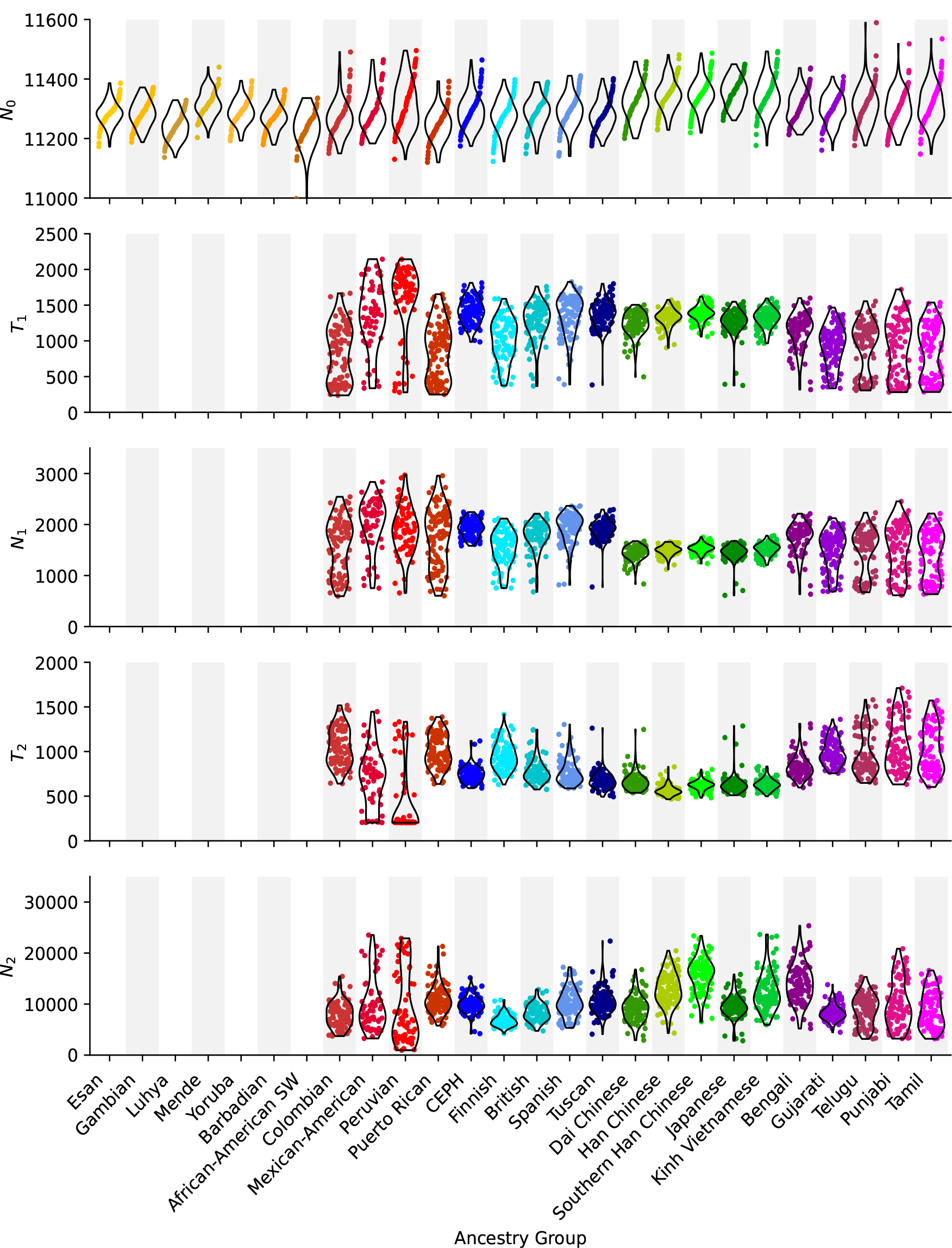
Parameter estimates for all individuals in the 1000 genomes data set. Within each ancestry group individuals are sorted by their estimate for the ancestral population size *N*_0_. The lack of correlation between the estimates for *N*_0_ and all other parameters lets the points in the bottom panels smear out. The mutation rate is assume to be *μ* = 2.36 *·* 10^−8^ per bp and generation, and the times *T*_1_ and *T*_2_ are measure in generations.

Where appropriate, i.e. for South American, European, and Asian populations, we also show estimates for the duration and population size during the bottleneck, *T*_1_ and *N*_1_, and the times and population sizes after the bottleneck, *T*_2_ and *N*_2_, respectively. We observe no significant correlation between our estimates for these quantities and the estimate for *N*_0_. The lack of correlation gives us confidence that our estimates of the population sizes at different times are independent of each other.

For European and Asian populations, all estimates are consistent and point to a bottleneck lasting roughly 1, 000 generations at a population size between 1, 000 and 2, 000 individuals, followed by population expansions to about 10, 000 individuals over the following 1, 000 generations. These estimates are close to what we know about the timing of the out-of-Africa bottleneck [31, 32]. Notably, the Peruvian ancestry group shows a significantly longer duration of the bottleneck period, offset by a shorter post-bottleneck period. This again is consistent with our knowledge of how humans migrated out of Africa and settled around the world. Some other individuals of South American ancestry show a similar pattern, although it is likely to be distorted by recent admixture events [7].

## 3 Discussion

In this article, we have studied the distribution of distances between heterozygous sites, or IBS tract lengths, in human genomes. We show that this distribution has a power-law tail with exponent −3. Using simple arguments and combining results for the distribution of times to the most recent common ancestor from coalescent theory and the stick-breaking process, we can analytically derive this powerlaw behavior.

Applied to a stationary population, we show that the length distribution of IBS tracts can be computed in closed form and depends on only two parameters: the size of the genome and *θ* = 4*Nμ*. This function fits very well to empirical data from genomes of African ancestry. Empirical data for populations that have undergone a bottleneck show significant deviations in the length distribution of IBS tracts. A generalization of our model to a demographic scenario with piecewise constant population sizes allows us to capture these deviations. We use these results to efficiently estimate population sizes before, during, and after the bottleneck, as well as the timing of the bottleneck for populations that migrated out of Africa. Interestingly, using our methodology, we do not see sufficient evidence to include a bottleneck in the model for African populations. This is in contrast to previous studies [6, 7], which inferred a mild bottleneck also for African individuals.

Some parts of the IBS tract length distributions are not well fitted by our model. Notably the upswing of the empirical distribution in the length range from 1 to 10 bp is not captured. This over-representation is due to multi-nucleotide substitutions, i.e. mutation events that change two nearby nucleotides at the same time. Such events are not accounted for in our model and generate many very short IBS tracts [33, 34]. Furthermore, it turns out that a large fraction of these events are also associated with small insertions and deletions in their neighborhood [34, 35], and we did not include such more complex changes of the DNA sequence in our modeling.

In addition, if we compare the empirical length distributions of IBS tracts for all individuals in a population (Figs. S3-S8), we observe that they are very consistent for lengths up to 100 kbp and become noisy beyond this length scale. This noise is due to the fact that only a few such tracts are observed in an individual’s genome, sometimes leading to over- or under-representation of very long IBS tracts. However, in certain populations, especially individuals of Gambian, Punjabi, Tamil, and Telugu ancestry, see Fig. S3(B) and S8(C-E), a large fraction of individuals show an upswing in the IBS tract length distribution for tracts longer than 100 kbp. This overrepresentation of very long IBS tracts might reflect inbreeding, which is more common in the above populations as previously reported [36]. The quantification of this effect could allow to define an inbreeding coefficient for single individuals without knowledge of a pedigree.

Here we only analyzed the distribution of heterozygous sites in humans. Of note, our method does not require a reference genome (except for the purpose of efficiently calling SNPs from short-read sequencing data) or the phase of SNPs along the maternal and paternal chromosomes. It is therefore straightforward to apply our methodology to other diploid species and haploid species if the genomes of two individuals are available. It can therefore be of great use in species where only little genomic information is available, for instance endangered or extinct species.

## 4 Materials and Methods

Files containing calls for phased variants against the hg38 reference genome of 2491 individuals have been downloaded from the website `ftp.1000genomes.ebi.ac.uk` of the 1000 genomes project [19, 20]. We excluded several individuals from the original 2504 individuals since they were reported to be children of other individuals in the same data set.

Insertion and deletions have been disregarded. The length of IBS tracts was computed as the difference *ℓ* = *p*_*i*+1_ − *p*_*i*_ of the positions of two consecutive heterozygous sites at *p*_*i*_ for on either of the parental chromosomes. We disregarded homozygous SNPs which appear on both chromosomes of an individual. Note that IBS tract of length *ℓ* has *ℓ* − 1 exactly matching base pairs in the two parental chromosomes. IBS tracts spanning over centromeres and at the very tips of chromosomes are disregarded as well. In our considerations the phase of a SNP is irrelevant.

The lengths distributions of all autosomes have been pooled after convincing ourselves that data from individual chromosomes do not show significant deviations from each other.

For graphical representation of count data in double-logarithmic length distributions, we have binned the data into bins of equal size along the logarithmic *x*-axis. We plot the density of counts on the *y*-axis, which is computed as the number of counts in an interval normalized by the length of the interval.

For graphs of the length distributions of IBS tracts for an ancestry group we aggregated the corresponding distributions for all individuals in that group and normalized the density by the number of individuals in that group.

All computations are performed using the Julia programming language [37]. The Laplace transform of the distribution of pairings *ñ*(*s, t*) was computed as described in the text, see Eq. (18). The second derivative, see Eq. (10), was computed using automatic differentiation [38].

To fit the model to empirical data we minimized the loss function

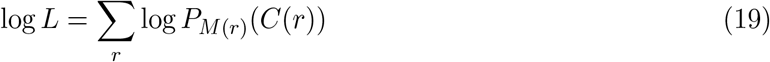

where *C*(*r*) is the count of IBS tracts of lengths *r* and *P*_*λ*_ denotes the Poisson distribution

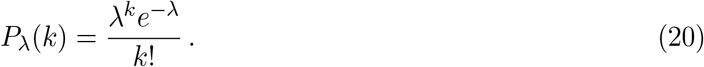

and lengths *r* ≤ 1, 000, 000 are considered. The modelled IBS tract length distribution *M* (*r*) depends on the sequence length, *L*, the initial population size, *N*_0_, as well as the lengths and population sizes, *T*_*i*_ and *N*_*i*_, of subsequent epochs, *i* = 1,…, *E*, see Eq. (18). The steady state model has *E* = 0.

We use Markov Chain Monte Carlo sampling to find the minimum of the loss function in the parameter space. Specifically, we use the No-U-Turn Sampler (NUTS) to adaptively set the path lengths in Hamiltonian Monte Carlo [39, 40]. The reported parameter values are mean values of at least 10 MCMC chains with 10, 000 samples after 10, 000 iterations to burn-in. We convinced ourselves that the MCMC chains converged. As an example we show the accumulative density distributions for all model parameters and the log likelihood for several chains in Fig. S9.

To access the stability of our parameter estimates and to compare models with different numbers of parameters (see Fig. 6) we considered 2, 000 bootstrap samples of IBS tract lengths and re-estimated all parameters.

## A Supplementary Material

### Normalisation of *M*

The length distribution of IBS tracts *M* is normalized, such that their total number is

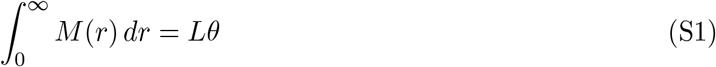

and their total length is

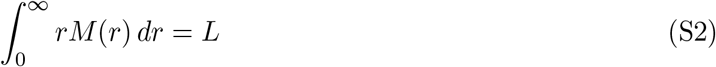

Therefore, the mean length of IBS tracts is *L/*(*Lθ*) = 1*/θ*. However, due to the power-law nature of the length distribution with exponent −3, the standard deviation of tract lengths diverges.

### The number of pairings for a population size increase

As described in the main text, for an instantaneous expansion of the population size at time *t*_*i*_, i.e. *N*_*i*+1_ *> N*_*i*_, we model that each present gamete will generate several offspring to make up the next generation. We assume that the number of offspring, *p*, for each gamete is distributed according to a truncated Poisson distribution:

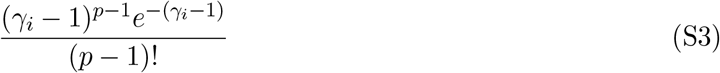

for *p ≥* 1 and vanishes for *p* = 0, i.e. we assume and that no gamete is lost. On average each gamete will have *γ*_*i*_ = *N*_*i*+1_*/N*_*i*_ *>* 1 offspring. A gamete has *p* offspring will generate *p*(*p* − 1)*/*2 pairs with *τ* = 0. For the whole population of 2*N*_*i*_ gametes the expected number of pairing with *τ* = 0 is

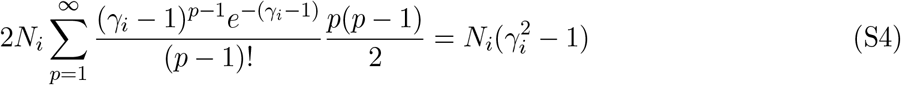

The resulting population at time 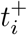 will be of size *N*_*i*+1_ and the number of pairings is

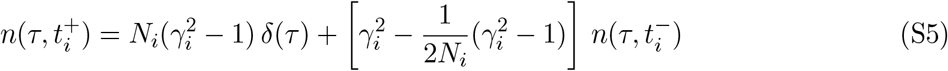

In Laplace space we recover Eq. (16).

### A model for an instantaneous bottleneck

Although the functions *M* (*r*) can be computed analytically the actual expression get quite long due to repeated applications of the transformations 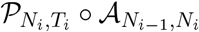, see Eq. (18), and especially due to the second derivative, see Eq. (10). However the function can easily be computed programmatically;especially for the second derivative automatic differentiation can be used, see Methods.

Here we only want to give the formula for *M* in a *E* = 2 epoch model with *N*_0_ *> N*_1_ *< N*_2_ assuming that the time spend in the bottleneck vanishes, i.e. *T*_1_ *→* 0. In this limit we have

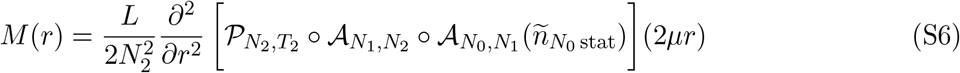

which can be computed to be

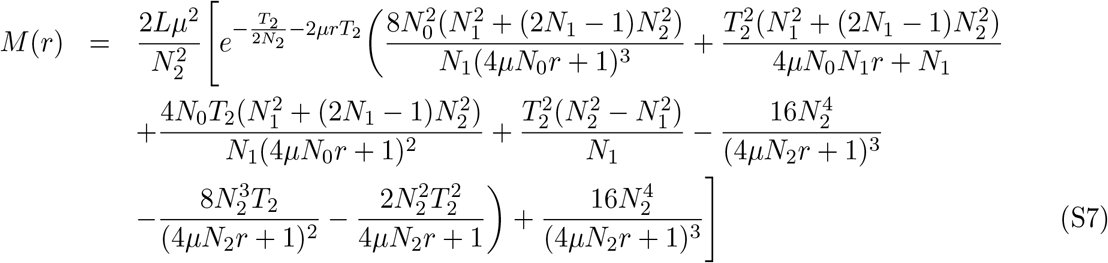

The humped part of this distribution is dominated by the term

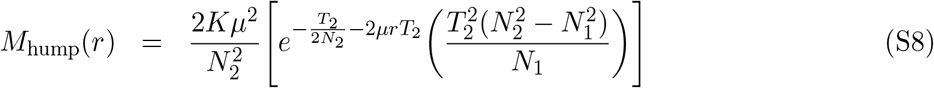

and the hump itself is located at *r*_hump_ = 3*/*(2*μT*_2_). Therefore older bottlenecks will be responsible for humps at smaller IBS tract lengths. Likewise, from the position of a hump one can roughly estimate the time of a bottleneck to be *T*_2_ = 3*/*2(*μr*_hump_). This general behavior will also hold for finite-time bottlenecks.

**Table S1:**
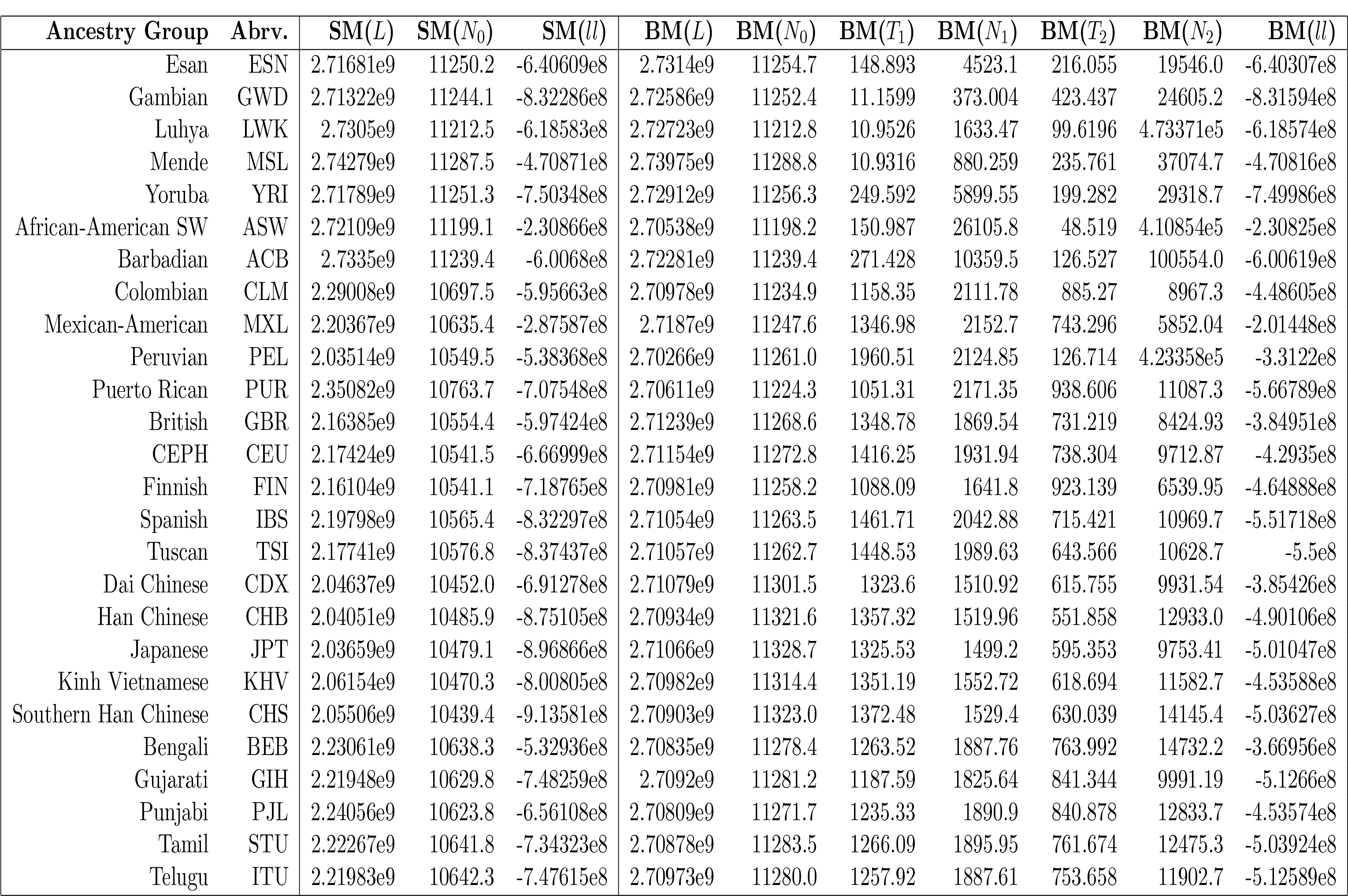
Parameter values for the stationary and the bottleneck model for different ancestry groups.

**Figure S1:**
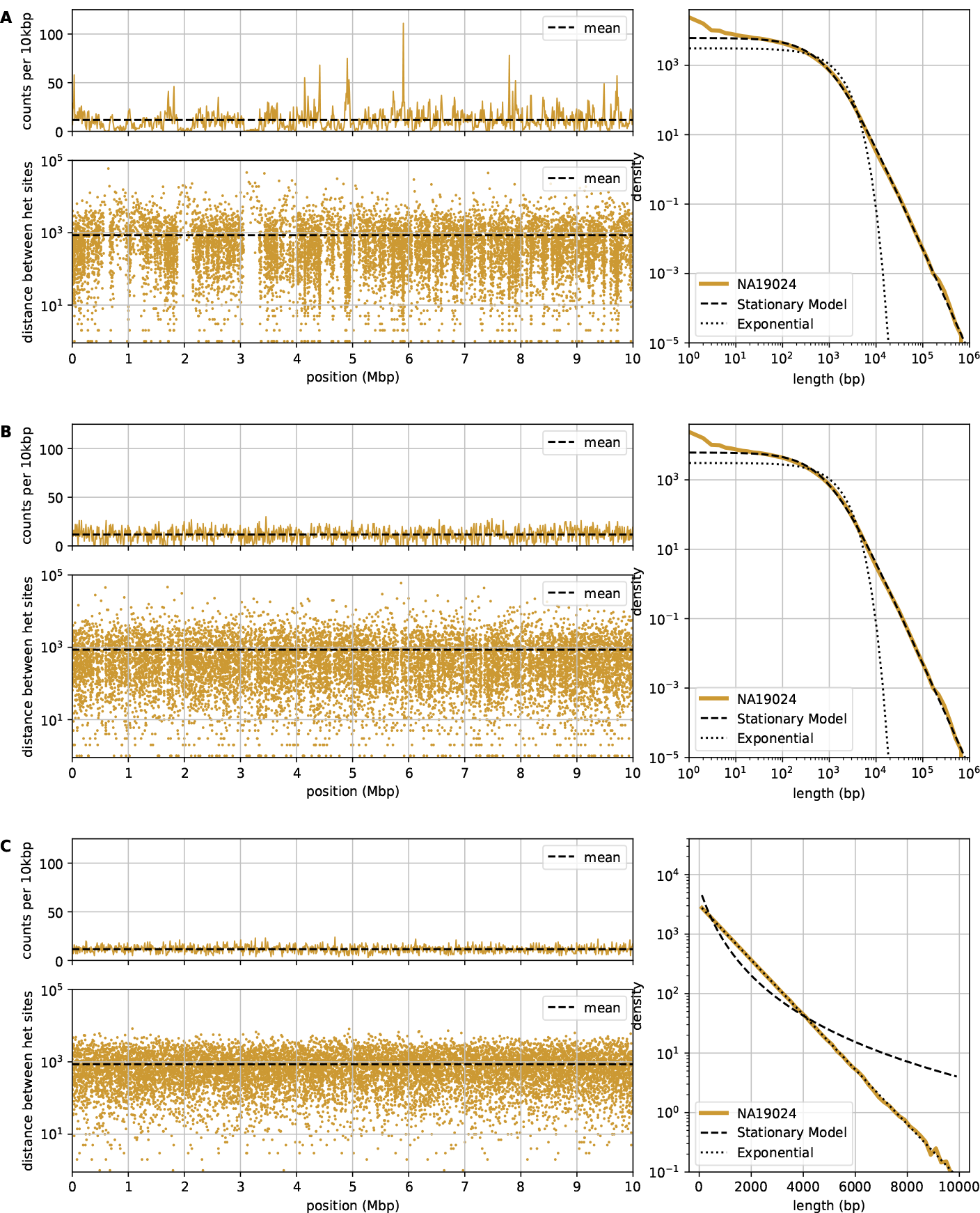
Comparison of the density and raindrop plot. (A) original data, (B) after shuffling the order of the IBS tracts, (C) after randomly re-sampling random positions of heterozygous sites. By construction all distributions have the same mean as indicated by the dashed line. The corresponding length distributions of IBS tracts are shown on the right. Note that these plots are on double logarithmic scale for (A) and (B) and on logarithmic/linear scale for (C).

**Figure S2:**
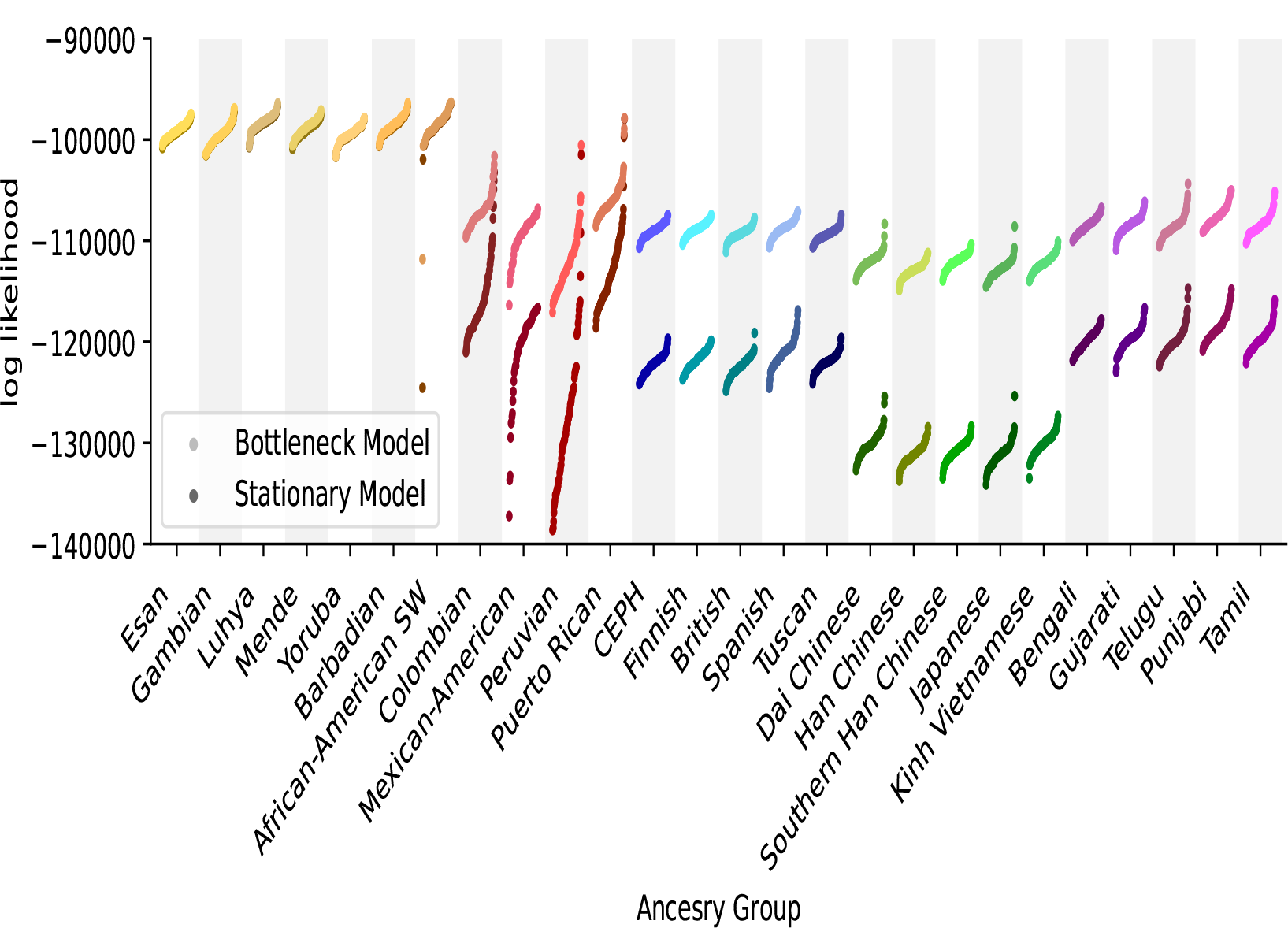
Likelihoods for fits with two different models for all individuals in the 1000 genomes dataset.

**Figure S3:**
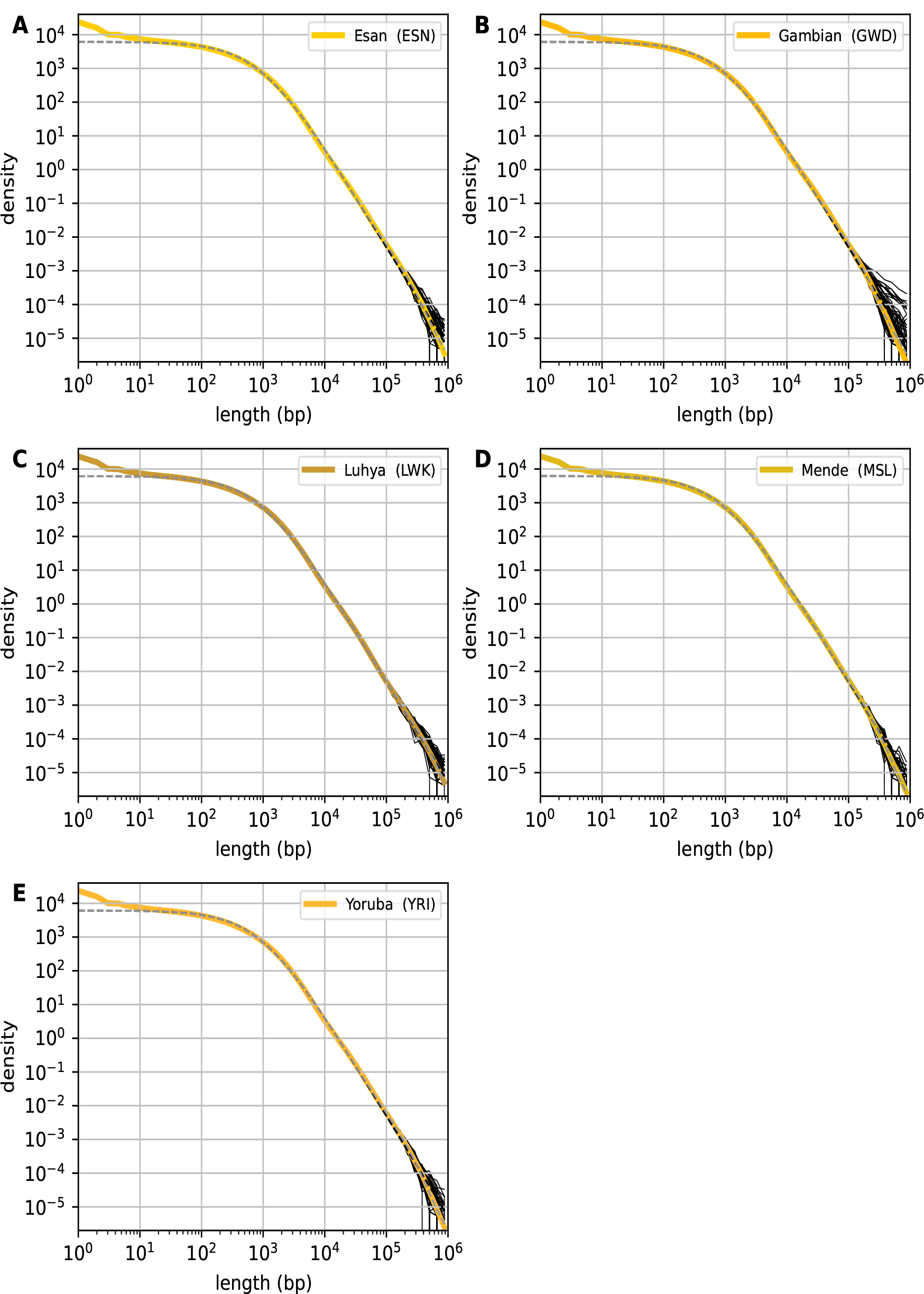
The length distributions of IBS tracts for all individuals in the African ancestry groups (black lines). The colored curve represents the group-specific mean length distribution. Fits using the stationary model (black dashed line) and the bottleneck model (gray dashed line) are included.

**Figure S4:**
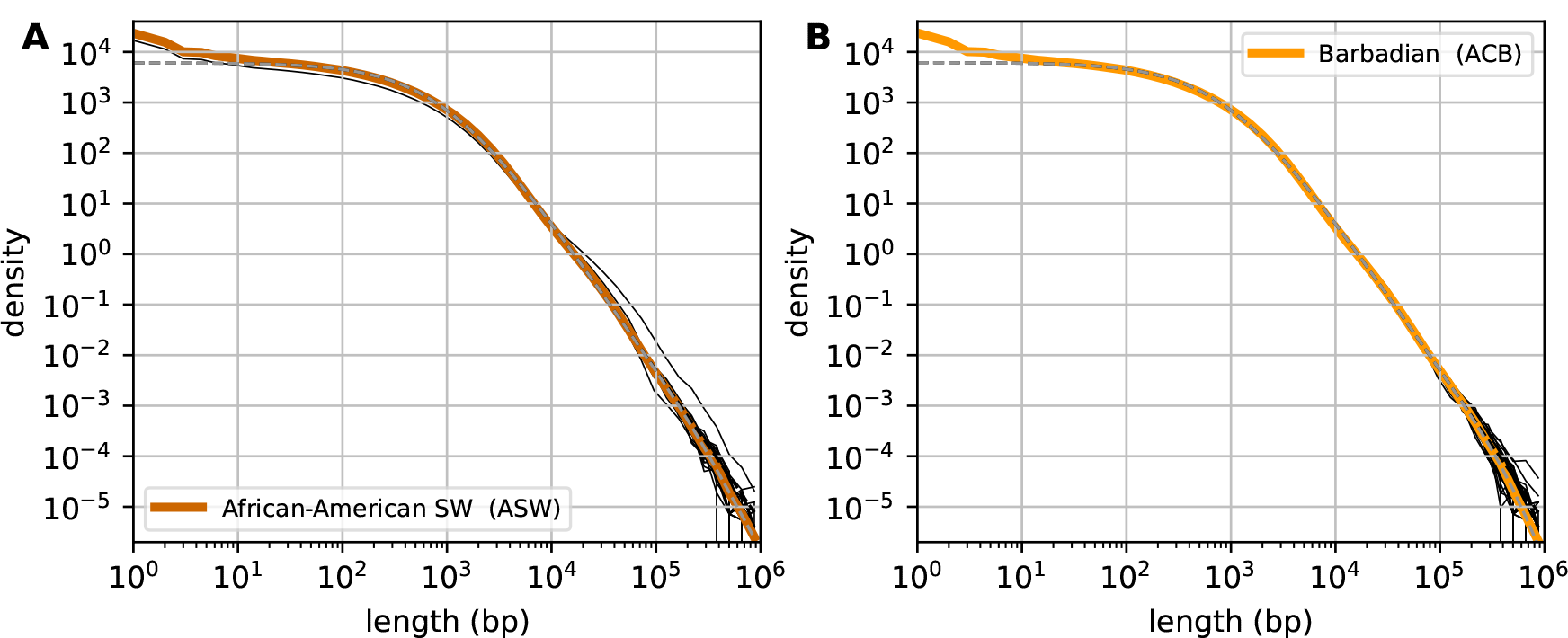
The length distributions of IBS tracts for all individuals in the African American ancestry groups (black lines). The colored curve represents the group-specific mean length distribution. Fits using the stationary model (black dashed line) and the bottleneck model (gray dashed line) are included.

**Figure S5:**
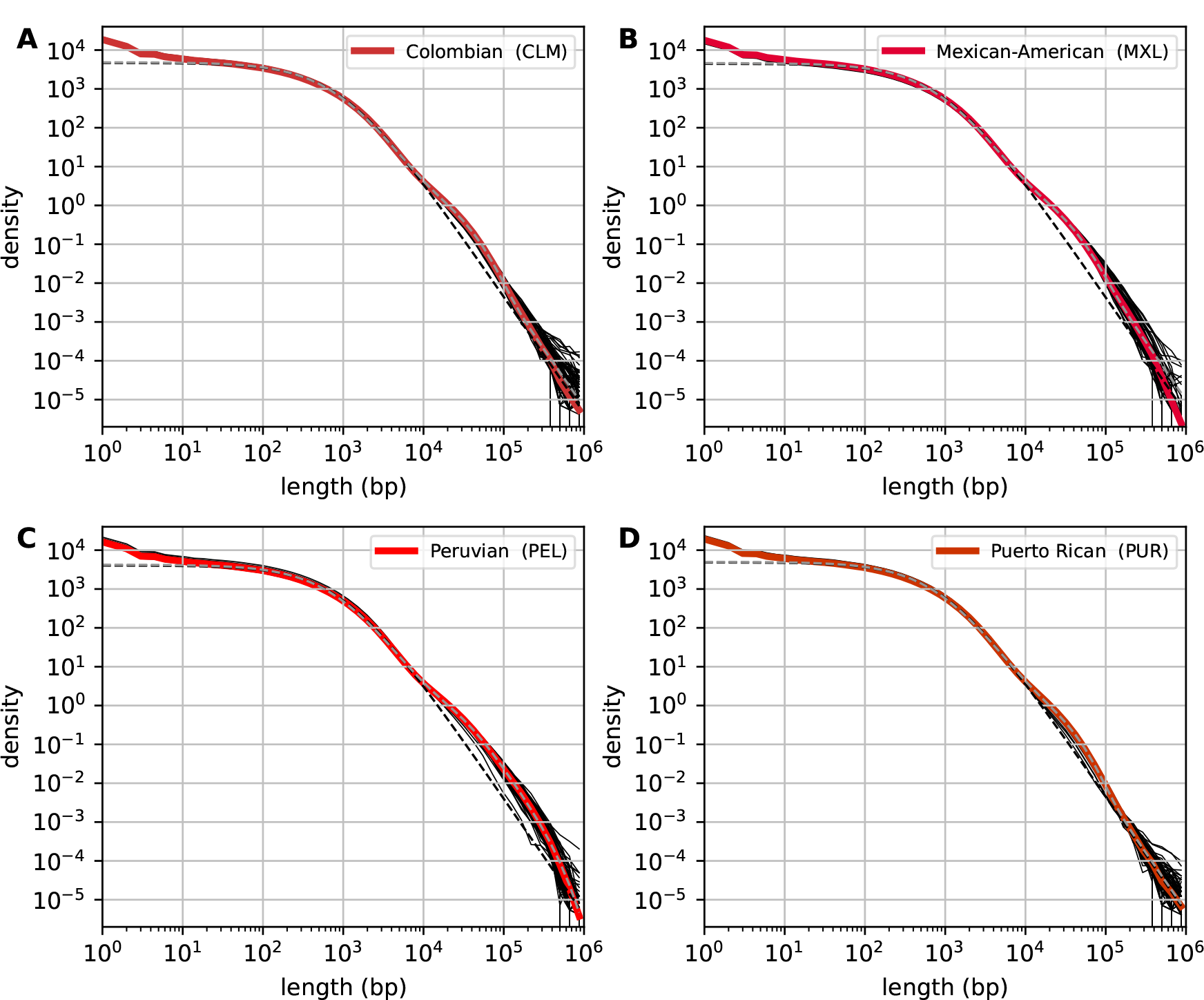
The length distributions of IBS tracts for all individuals in the South American ancestry groups (black lines). The colored curve represents the group-specific mean length distribution. Fits using the stationary model (black dashed line) and the bottleneck model (gray dashed line) are included.

**Figure S6:**
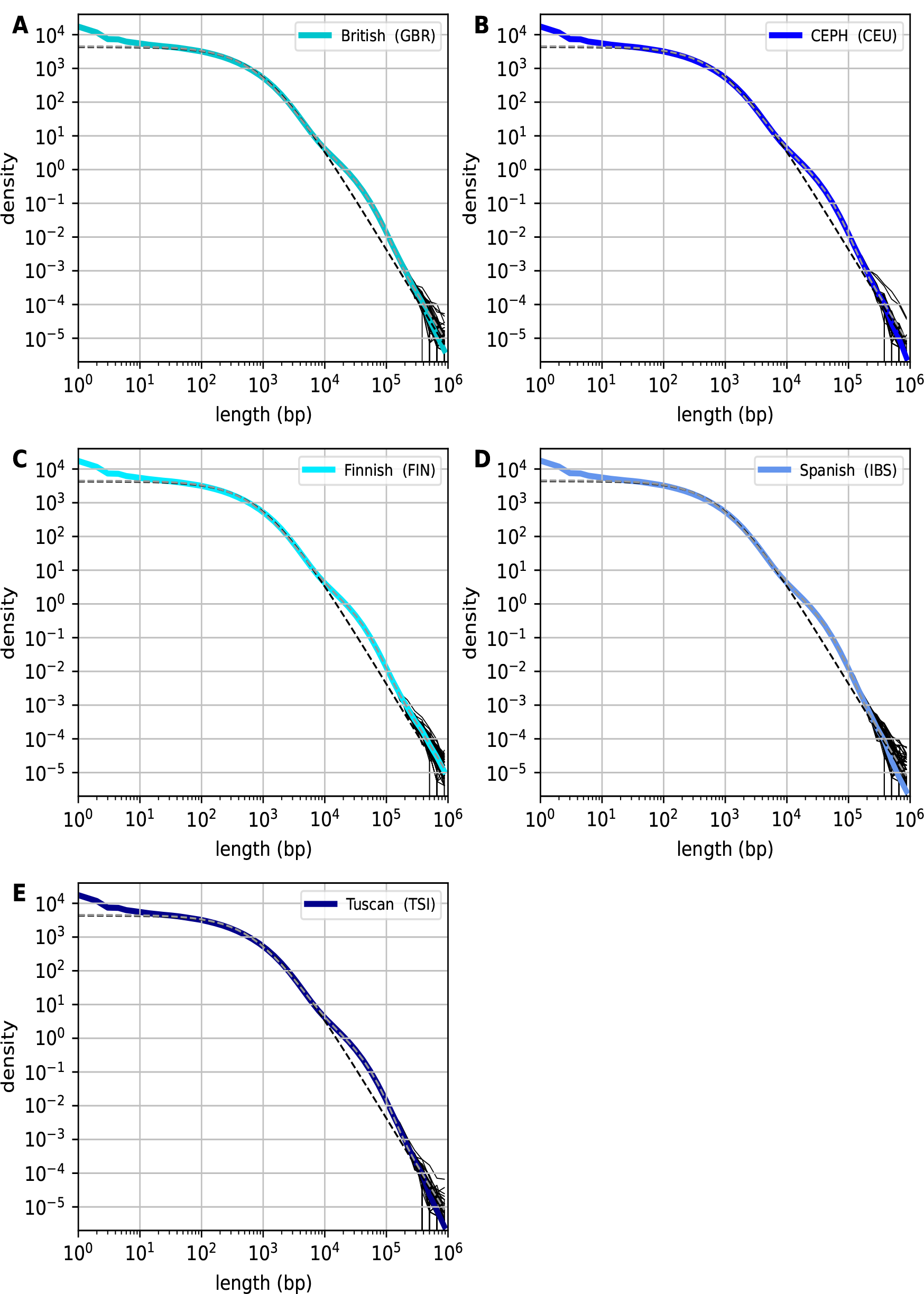
The length distributions of IBS tracts for all individuals in the European ancestry groups (black lines). The colored curve represents the group-specific mean length distribution. Fits using the stationary model (black dashed line) and the bottleneck model (gray dashed line) are included.

**Figure S7:**
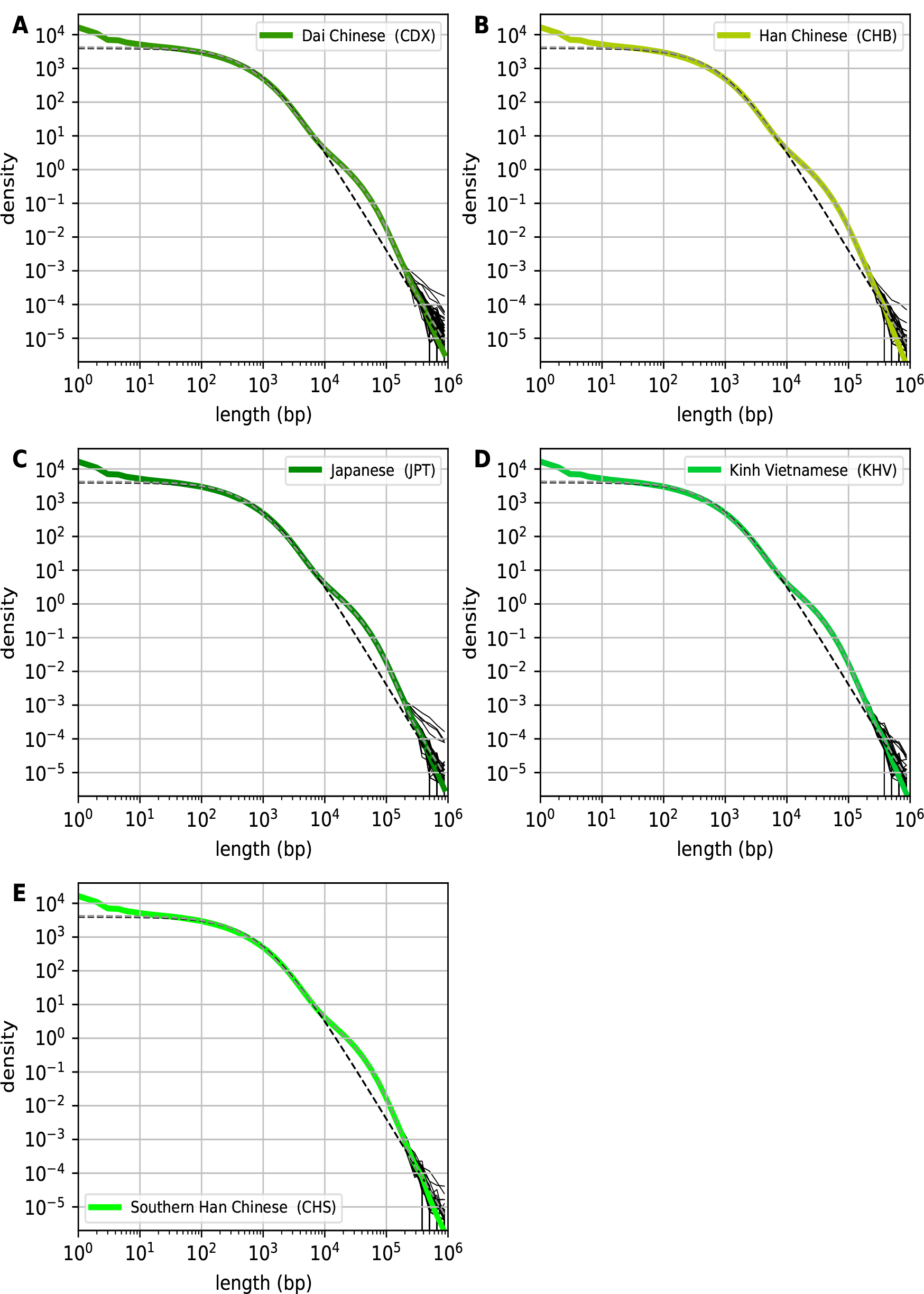
The length distributions of IBS tracts for all individuals in the East Asian ancestry groups (black lines). The colored curve represents the group-specific mean length distribution. Fits using the stationary model (black dashed line) and the bottleneck model (gray dashed line) are included.

**Figure S8:**
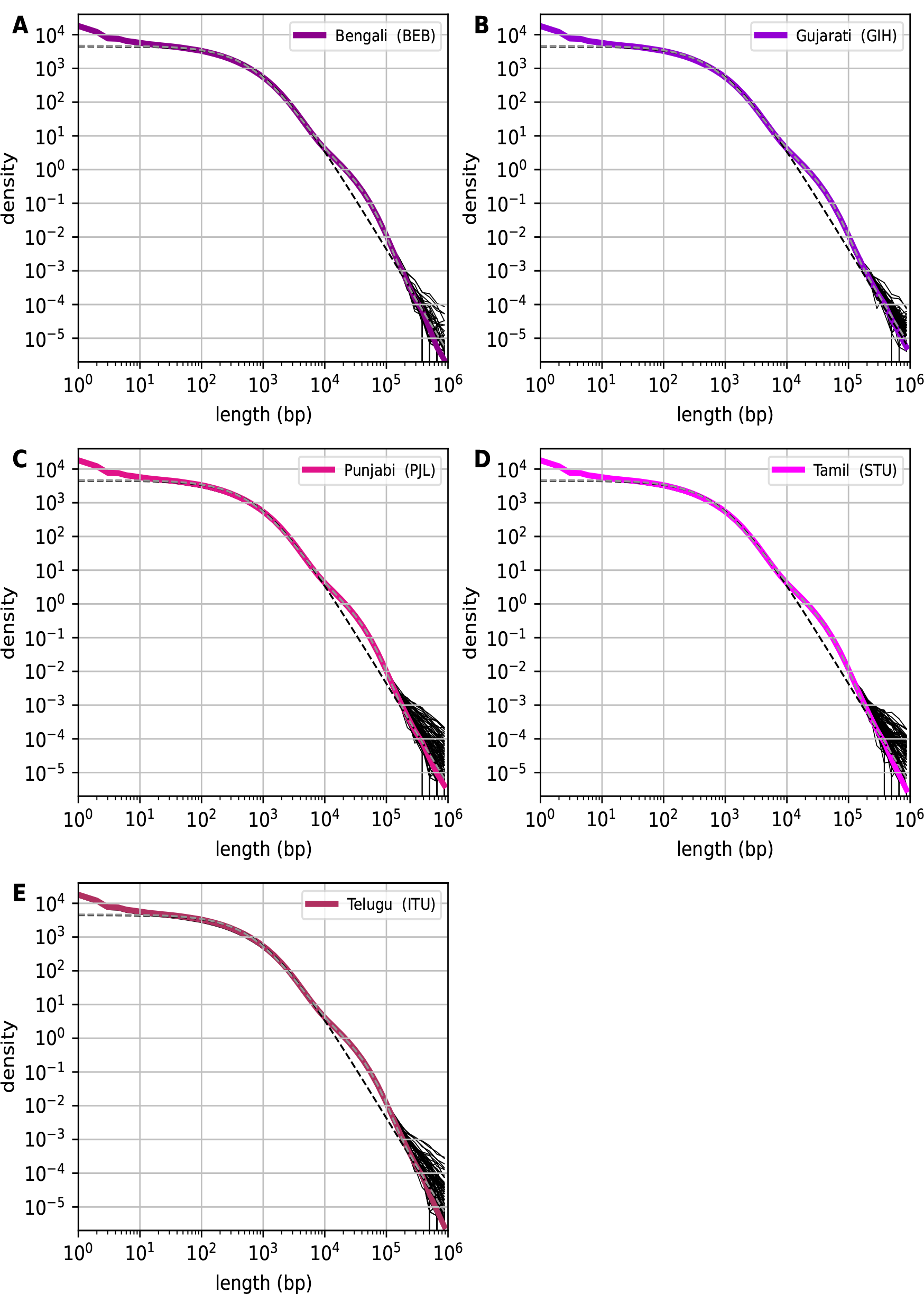
The length distributions of IBS tracts for all individuals in the South Asian ancestry groups (black lines). The colored curve represents the group-specific mean length distribution. Fits using the stationary model (black dashed line) and the bottleneck model (gray dashed line) are included.

**Figure S9:**
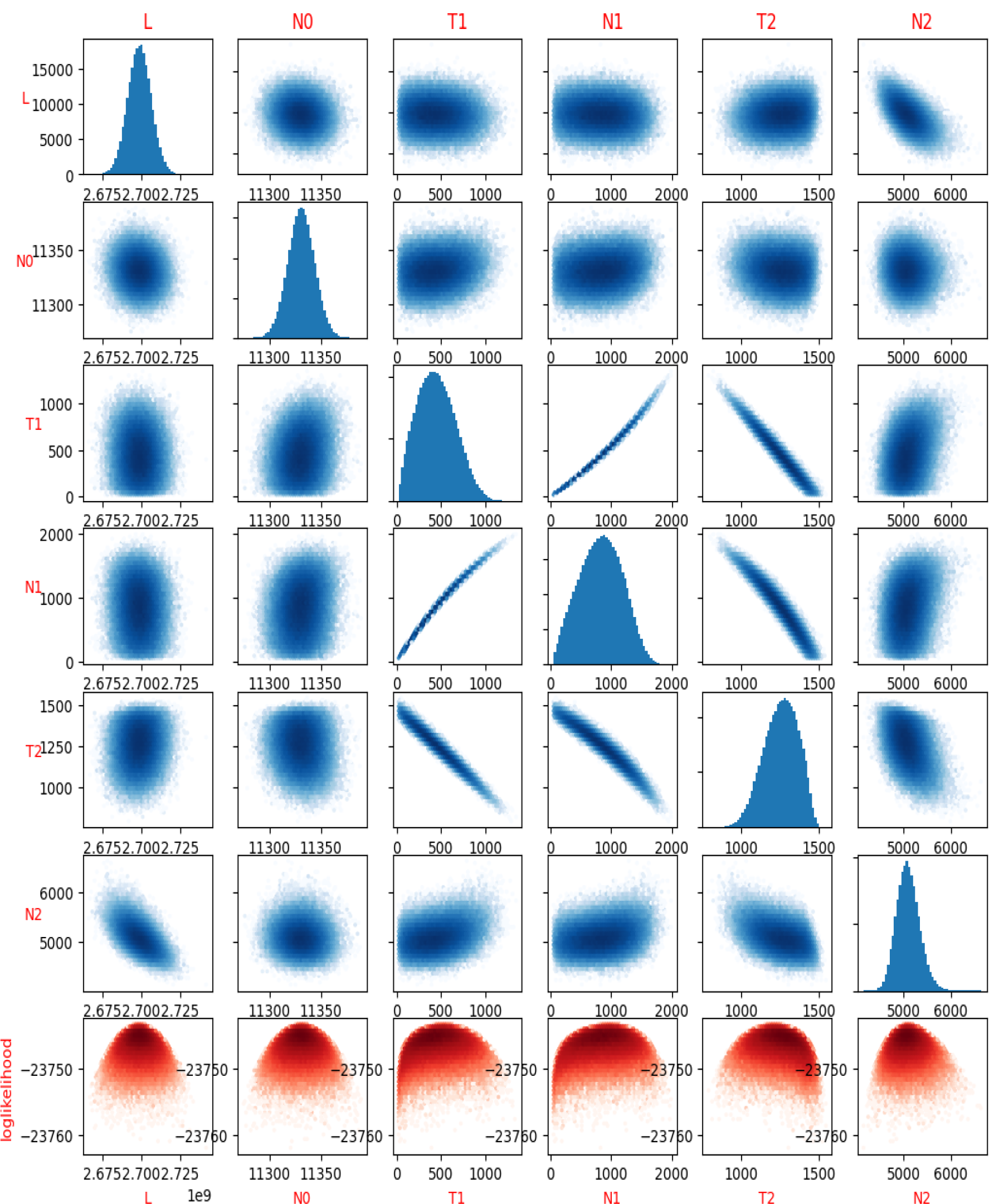
Example of an scatter density plot of 6 model parameters and the loglikelihood of 10 MCMC chains to fit a bottleneck model.

## Notes

### Competing Interest Statement

The authors have declared no competing interest.

## References

[1] The International SNP Map Working Group, Cold Spring Harbor Laboratories: Ravi Sachidanandam, et al. “A map of human genome sequence variation containing 1.42 million single nucleotide polymorphisms”. In: Nature 409.6822 (2001), pp. 928–933. ISSN: 0028-0836. doi: 10.1038/35057149.

[2] Wayne S. Kendal. “An Exponential Dispersion Model for the Distribution of Human Single Nucleotide Polymorphisms”. In: Molecular Biology and Evolution 20.4 (2003), pp. 579–590. ISSN: 0737-4038. doi: 10.1093/molbev/msg057.

[3] Bernhard Haubold, Nora Pierstorff, Friedrich Möller, and Thomas Wiehe. “Genome comparison without alignment using shortest unique substrings.” In: BMC bioinformatics 6.1 (2004), p. 123. doi: 10.1186/1471-2105-6-123.

[4] Bernhard Haubold and Peter Pfaffelhuber. “Alignment-Free Population Genomics: An Efficient Estimator of Sequence Diversity”. In: G3: Genes—Genomes—Genetics 2.8 (2012), pp. 883–889. doi: 10.1534/g3.112.002527.

[5] Gilean A.T McVean and Niall J Cardin. “Approximating the coalescent with recombination”. In: Philosophical Transactions of the Royal Society B: Biological Sciences 360.1459 (2005), pp. 1387–1393. ISSN: 0962-8436. doi: 10.1098/rstb.2005.1673.

[6] Heng Li and Richard Durbin. “Inference of human population history from individual wholegenome sequences.” English. In: Nature 475.7357 (July 2011), pp. 493–496. doi: 10.1038/nature10231.

[7] Stephan Schiffels and Richard Durbin. “Inferring human population size and separation history from multiple genome sequences”. In: Nature genetics 46.8 (2014), pp. 919–925. ISSN: 1061-4036. doi: 10.1038/ng.3015.

[8] Jonathan Terhorst, John A Kamm, and Yun S Song. “Robust and scalable inference of population history from hundreds of unphased whole genomes”. In: Nature Genetics 49.2 (2017), pp. 303–309. ISSN: 1061-4036. doi: 10.1038/ng.3748.

[9] Pier Francesco Palamara, Jonathan Terhorst, Yun S. Song, and Alkes L. Price. “High-throughput inference of pairwise coalescence times identifies signals of selection and enriched disease heritability”. In: Nature Genetics 50.9 (2018), pp. 1311–1317. ISSN: 1061-4036. doi: 10.1038/s41588-018-0177-x.

[10] Thibaut Paul Patrick Sellinger, Diala Abu Awad, Markus Moest, and Aurélien Tellier. “Inference of past demography, dormancy and self-fertilization rates from whole genome sequence data”. In: PLoS Genetics 16.4 (2020), e1008698. ISSN: 1553-7390. doi: 10.1371/journal.pgen.1008698.

[11] Junfeng Liu, Xianchao Ji, and Hua Chen. “Beta-PSMC: uncovering more detailed population history using beta distribution”. In: BMC Genomics 23.1 (2022), p. 785. doi: 10.1186/s12864-022-09021-6.

[12] Simon Gravel, Brenna M. Henn, Ryan N. Gutenkunst, et al. “Demographic history and rare allele sharing among human populations”. In: Proceedings of the National Academy of Sciences 108.29 (2011), pp. 11983–11988. ISSN: 0027-8424. doi: 10.1073/pnas.1019276108.

[13] Thibaut Paul Patrick Sellinger, Diala Abu-Awad, and Aurélien Tellier. “Limits and convergence properties of the sequentially Markovian coalescent.” In: Molecular ecology resources 21.7 (2020), pp. 2231–2248. ISSN: 1755-098X. doi: 10.1111/1755-0998.13416.

[14] Kelley Harris and Rasmus Nielsen. “Inferring Demographic History from a Spectrum of Shared Haplotype Lengths”. In: PLoS Genetics 9.6 (2013), e1003521. ISSN: 1553-7390. doi: 10.1371/journal.pgen.1003521.

[15] Eric S. Lander and David Botstein. “Homozygosity Mapping: A Way to Map Human Recessive Traits with the DNA of Inbred Children”. In: Science 236.4808 (1987), pp. 1567–1570. ISSN: 0036-8075. doi: 10.1126/science.2884728.

[16] Karl W. Broman and James L. Weber. “Long Homozygous Chromosomal Segments in Reference Families from the Centre d’Étude du Polymorphisme Humain”. In: The American Journal of Human Genetics 65.6 (1999), pp. 1493–1500. ISSN: 0002-9297. doi: 10.1086/302661.

[17] Kelly A. Frazer, Dennis G. Ballinger, David R. Cox, et al. “A second generation human haplotype map of over 3.1 million SNPs”. In: Nature 449.7164 (2007), pp. 851–861. ISSN: 0028-0836. doi: 10.1038/nature06258.

[18] Francisco C. Ceballos, Peter K. Joshi, David W. Clark, Michéle Ramsay, and James F. Wilson. “Runs of homozygosity: windows into population history and trait architecture”. In: Nature Reviews Genetics 19.4 (2018), pp. 220–234. ISSN: 1471-0056. doi: 10.1038/nrg.2017.109.

[19] Adam Auton, Gonçalo R. Abecasis, David M. Altshuler, et al. “A global reference for human genetic variation”. In: Nature 526.7571 (2015), pp. 68–74. ISSN: 0028-0836. doi: 10.1038/nature15393.

[20] Marta Byrska-Bishop, Uday S. Evani, Xuefang Zhao, et al. “High-coverage whole-genome sequencing of the expanded 1000 Genomes Project cohort including 602 trios”. In: Cell 185.18 (2022), 3426–3440.e19. ISSN: 0092-8674. doi: 10.1016/j.cell.2022.08.004.

[21] Motoo Kimura. “The number of heterozygous nucleotide sites maintained in a finite population due to steady flux of mutations”. In: Genetics 61.4 (1969), pp. 893–903. ISSN: 1943-2631. doi: 10.1093/genetics/61.4.893.

[22] G A Watterson. “On the number of segregating sites in genetical models without recombination.” English. In: Theoretical population biology 7.2 (1975), pp. 256–276. doi: 10.1016/0040-5809(75)90020-9.

[23] W.J. Ewens. “A note on the sampling theory for infinite alleles and infinite sites models”. In: Theoretical Population Biology 6.2 (1974), pp. 143–148. ISSN: 0040-5809. doi: 10.1016/0040-5809(74)90020-3.

[24] J. Hein, M.H. Schierup, and C. Wiuf. Gene Genealogies, Variation and Evolution: A Primer in Coalescent Theory. Oxford University Press. ISBN: 9780198529965.

[25] Serena Nik-Zainal, Ludmil B. Alexandrov, David C. Wedge, et al. “Mutational Processes Molding the Genomes of 21 Breast Cancers”. In: Cell 149.5 (2012), pp. 979–993. ISSN: 0092-8674. doi: 10.1016/j.cell.2012.04.024.

[26] Warren J. Ewens. Mathematical Population Genetics. New York: Springer, 2004. ISBN: 1441918981. doi: 10.1007/978-0-387-21822-9.

[27] Florian Massip and Peter F Arndt. “Neutral Evolution of Duplicated DNA: An Evolutionary Stick-Breaking Process Causes Scale-Invariant Behavior”. English. In: Physical Review Letters 110.14 (2013), p. 148101. doi: 10.1103/physrevlett.110.148101. URL: http://prl.aps.org/abstract/PRL/v110/i14/e148101.

[28] Peter F Arndt. “Sequential and continuous time stick-breaking”. In: Journal of Statistical Mechanics: Theory and Experiment 2019.6 (June 2019), pp. 064003–9. doi: 10.1088/1742-5468/ab1dd8.

[29] Michael Sheinman, Florian Massip, and Peter F Arndt. “Statistical Properties of Pairwise Distances between Leaves on a Random Yule Tree.” English. In: PloS one 10.3 (2015), e0120206. doi: 10.1371/journal.pone.0120206.

[30] Florian Massip, Michael Sheinman, Sophie Schbath, and Peter F Arndt. “How evolution of genomes is reflected in exact DNA sequence match statistics.” English. In: Molecular biology and evolution 32.2 (2015), pp. 524–535. doi: 10.1093/molbev/msu313.

[31] Paul Mellars. “Going East: New Genetic and Archaeological Perspectives on the Modern Human Colonization of Eurasia”. In: Science 313.5788 (2006), pp. 796–800. ISSN: 0036-8075. doi: 10.1126/science.1128402.

[32] Brenna M. Henn, L. L. Cavalli-Sforza, and Marcus W. Feldman. “The great human expansion”. In: Proceedings of the National Academy of Sciences 109.44 (2012), pp. 17758–17764. ISSN: 0027-8424. doi: 10.1073/pnas.1212380109.

[33] Jeffrey A. Rosenfeld, Anil K. Malhotra, and Todd Lencz. “Novel multi-nucleotide polymorphisms in the human genome characterized by whole genome and exome sequencing”. In: Nucleic Acids Research 38.18 (2010), pp. 6102–6111. ISSN: 0305-1048. doi: 10.1093/nar/gkq408.

[34] Qingbo Wang, Emma Pierce-Hoffman, Beryl B Cummings, et al. “Landscape of multi-nucleotide variants in 125,748 human exomes and 15,708 genomes”. In: Nature Communications 11.1 (2020), p. 2539. doi: 10.1038/s41467-019-12438-5.

[35] Tsung-Yu Lu, The Human Genome Structural Variation Consortium, Katherine M Munson, et al. “Profiling variable-number tandem repeat variation across populations using repeat-pangenome graphs”. In: Nature Communications 12.1 (2021), p. 4250. doi: 10.1038/s41467-021-24378-0.

[36] Steven Gazal, Mourad Sahbatou, Marie-Claude Babron, Emmanuelle Génin, and Anne-Louise Leutenegger. “High level of inbreeding in final phase of 1000 Genomes Project”. In: Scientific Reports 5.1 (2015), p. 17453. doi: 10.1038/srep17453.

[37] Jeff Bezanson, Alan Edelman, Stefan Karpinski, and Viral B Shah. “Julia: A Fresh Approach to Numerical Computing”. In: SIAM Review 59.1 (2017), pp. 65–98. ISSN: 0036-1445. doi: 10.1137/141000671.

[38] Jarrett Revels, Miles Lubin, and Theodore Papamarkou. “Forward-Mode Automatic Differentiation in Julia”. In: arXiv (2016). doi: 10.48550/arxiv.1607.07892. eprint: 1607.07892.

[39] Matthew D Hoffman and Andrew Gelman. “The No-U-Turn Sampler: Adaptively Setting Path Lengths in Hamiltonian Monte Carlo”. In: arXiv (2011). doi: 10.48550/arxiv.1111.4246. eprint: 1111.4246.

[40] Hong Ge, Kai Xu, and Zoubin Ghahramani. “Turing: A Language for Flexible Probabilistic Inference”. In: Proceedings of Machine Learning Research 84 (2018). Ed. by [“Storkey, {Perez-Cruz, Amos and}, and Fernando”], pp. 1682–1690. URL: https://proceedings.mlr.press/v84/ge18b.html.

